# The dorsolateral striatum regulates habits by way of performance vigor when actions are initiated

**DOI:** 10.1101/426031

**Authors:** Adam C.G. Crego, Fabián Štoček, Alec G. Marchuk, James E. Carmichael, Matthijs A.A. van der Meer, Kyle S. Smith

**Affiliations:** Department of Psychological and Brain Sciences, Dartmouth College, Hanover, New Hampshire 03755

## Abstract

Despite clear evidence linking basal ganglia control to habits, it remains unclear through what mechanisms this control occurs. Here, we demonstrate that a key function of the dorsolateral striatum (DLS) is to regulate the vigor of a learned behavior at the moment of initiation in a habit-promoting manner. Shifts in vigor by phasic DLS perturbations coincide closely with how outcome-insensitive (i.e., habitual) the behaviors are in both response-based and cue-based task situations. Surprisingly, the control over habit strength by way of changes in vigor occurs without consistent changes in accuracy, suggesting that mechanisms controlling habit and vigor are dissociable from performance governed task rules. Finally, we show that increased DLS activity improves vigor preferentially when learned outcome values are stable, while reduced DLS activity dampens vigor preferentially when outcome values change. These data indicate that improving action vigor could be a principle route by which the basal ganglia facilitate habits.

## Introduction

Habits are behavioral routines characterized by their automaticity, consistency, and resistance to change. A key node within the habit-promoting network is thought to be the dorsolateral striatum (DLS, primate putamen homologue), as DLS dysfunction results in behaviors that are less habitual^1,2^. Reduced habit strength can be detected in poorly sequenced actions, the use of place-based rules instead of response-based rules in maze tasks, and a sensitivity of behavior to changes in the relationship between a learned response and its outcome in instrumental tasks^3-9^. A separate line of research has implicated the basal ganglia in controlling performance vigor^10-12^, namely that it aids in pushing ongoing movement kinematics^13^ by affecting the speed of an action (i.e., vigor)^10^. This function would be analogous to swinging on a swing, with the assistance of someone pushing one forward (i.e., the DLS) affecting the vigor of this ongoing action. To date, it is generally unclear how measures of vigor relate to formal measures of habits in behavior or in ongoing basal ganglia activity. One entry point to studying this issue, used here, is a prominent activity signal in the DLS that occurs during learned habits and skills. Specifically, there is often a burst of DLS activity (200-500 ms) at the initiation of a habit or a well-learned action sequence^14-18^. This signaling event has been found in a range of species and DLS neuronal subtypes^14,16,19^, and can also occur in wider brain networks^17,18,20-23^. Notably, increases in the magnitude of action-start-related activity in the DLS and its substantia nigra pars compacta (SNc) input are closely related to behavioral changes reflective of increased vigor^12,17,18^. Within the DLS in particular, the activity is predictive at the single trial level of reduced performance duration and reduced deliberations (vicarious trial-and-error (VTE) head movements) at decision points^18^. Action-start DLS activity has also been found to be related to a defining characteristic of habits, outcome-insensitivity, which develops in parallel to the above vigor improvements; yet this relation is at the session level rather than trial-to-trial. Two hypotheses are raised and tested here: (1) the action-initiation time window is critical for DLS control over habits; (2) improvements in vigor are generated by this signal and lead to an increase in habit strength reflected in outcome insensitivity.

## Results

Rats were first trained on a DLS-dependent plus-maze response task^24,25^, in which a specific turn (e.g., turn left) at a choice point was always paired with an appetitive reward regardless of starting location (Fig. 1A). Training proceeded until 3 consecutive days of ≥75% accuracy was achieved (Fig. 1E). A series of test days followed during which the DLS was optogenetically stimulated (channelrhodopsin; ChR2) or inhibited (halorhodopsin; eNpHR3.0) for 0.5 sec at the onset of the maze runs, for 0.5 sec during the middle (maze center) of the runs, or on a 0.5 sec on/off cycle for the duration of the runs. These manipulations were compared to a sham control condition, in which animals were treated identically but lacked DLS opsin expression (Fig. 1C). We non-specifically targeted DLS neurons given that a majority of projection neurons and interneurons are active at behavior onset^l6,l8,19,26^ and play a critical role in habits^27-30^. Confocal imaging (GFP immunostain/neurotrace) revealed roughly an estimated 83% of DLS neurons expressed the viral molecules (Fig. 1B). Separate awake/behaving recordings confirmed ChR2-mediated stimulation efficacy (Fig. 1D); similar eNpHR3.0-mediated inhibition has been reported^31-33^.

**Figure 1.**
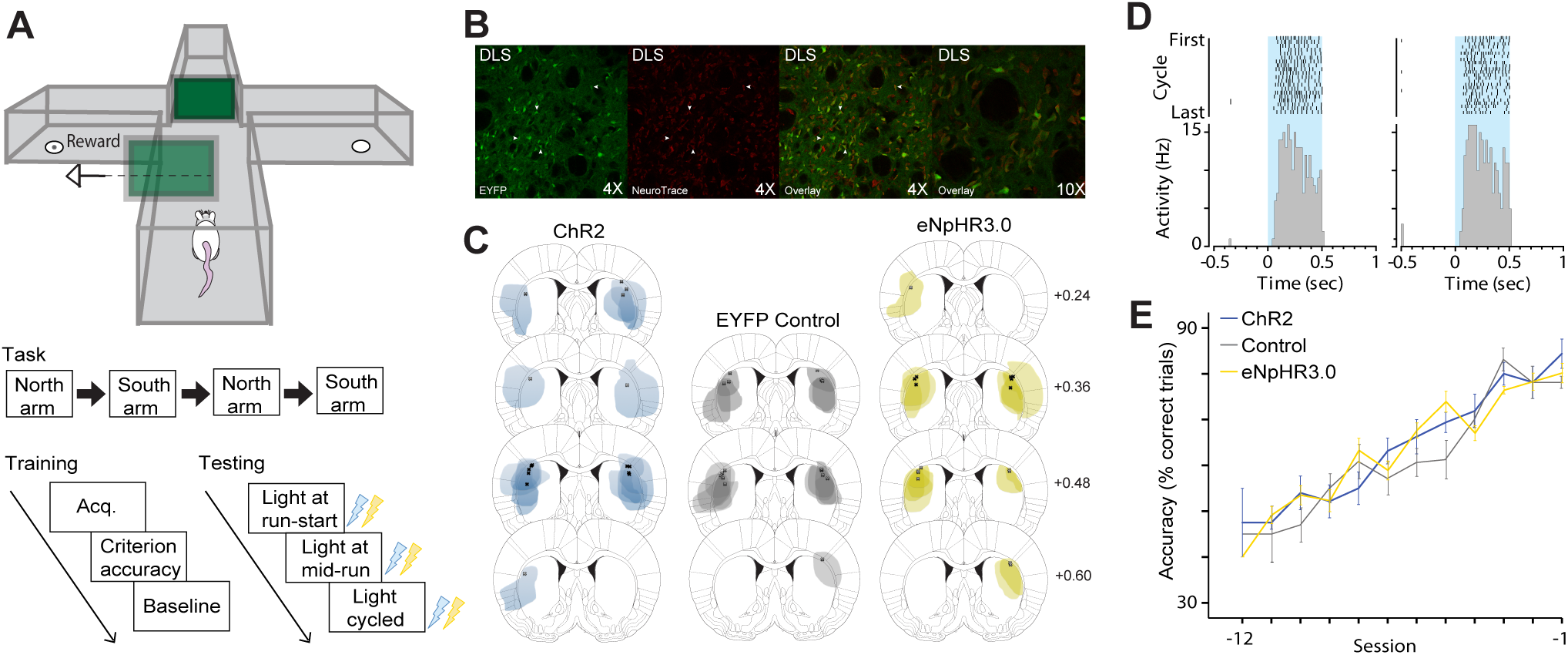
Response task. **(A)** Top, maze task schematic illustrating left turn example (green = gates). Middle, example protocol of trial blocks with start locations for a session. Bottom left, session order for task acquisition (through 3 days of ≥75% accuracy), and baseline test (fiber cables connected but no light). Bottom right, example DLS manipulation protocol (0.5 sec light delivery) on test days. **(B)** Confocal images of DLS section from left to right: EYFP, Neurotrace, overlay low-mag, overlay high-mag. White arrows show same neuron examples across labeling. **(C)** Expression areas of ChR2-EYFP, EYFP control, and eNpHR3.0-EYFP across animals. Semi-transparent shading denotes expression per animal; shading overlaid across animals. Numbers denote anteroposterior plane in mm relative to bregma. **(D)** Example rasters and histograms (0.02 sec/bin) of a single DLS unit putatively expressing ChR2 recorded under anesthesia. Left, firing excitation to 0.5 sec light delivery (blue shading) given every 1 min, approximating the run-start or mid-run task illuminations. Right, same unit showing firing excitation to 0.5 sec light delivery at a 0.5 sec on/off cycle, approximating the cycled task illuminations. See Supplementary Fig. 1 for all responsive units. **(E)** Run accuracy in sessions leading up to the final day of criterion performance, separated by ChR2 (blue), control (gray), and eNpHR3.0 (yellow) groups. No main effect of Group (*F*_2,243_=1.558, *P*=0.213) or Group/Day interaction (*F*_22,243_=1.349, *P*=0.144), but main effect of Day (*F*_11,243_=44.861, *P*<0.001). For all panels, lines and errors show mean ± SEM. ChR2, channelrhodopsin-2; eNpHR3.0, halorhodopsin. cc, corpus collosum; DLS, dorsolateral striatum.

Phasically activating the DLS at run-start resulted in increased running vigor by several behavioral indices. The cumulative duration of runs and the deliberative movements (VTEs) at the choice point in the center of the maze were markedly reduced compared to controls or to animals with run-start inhibition (Fig. 2C, 2B). The standard deviation of the cumulative duration was also reduced, indicating greater trial-to-trial consistency in runs (Fig. 2D). In stark contrast, mid-run stimulation had no effect on any of these vigor measures in the same animals, which is a similar effect to a recent report on dopamine neurons^12^. This is notable given that this mid-run time period is when deliberations would occur. Cycled on/off stimulation (which included a 0.5 sec pulse at start) produced similar effects to run-start stimulation, suggesting that the run-start time carried this effect. Analysis of run durations in the major maze segments (start-arm, maze-center, endarm; Fig. 2E; Supplementary Table 1) indicated that changes in vigor were not related strictly to changes in the initiation of the run when DLS perturbation occurred. Phasic DLS inhibition was effective at modulating vigor too, but less robustly so. Run-start inhibition itself did not substantially increase run duration compared to controls, but run-center inhibition did moderately and cycled inhibition did robustly. Deliberations were increased by cycled inhibition as well (Fig 2B; Supplementary Table 1).

**Figure 2.**
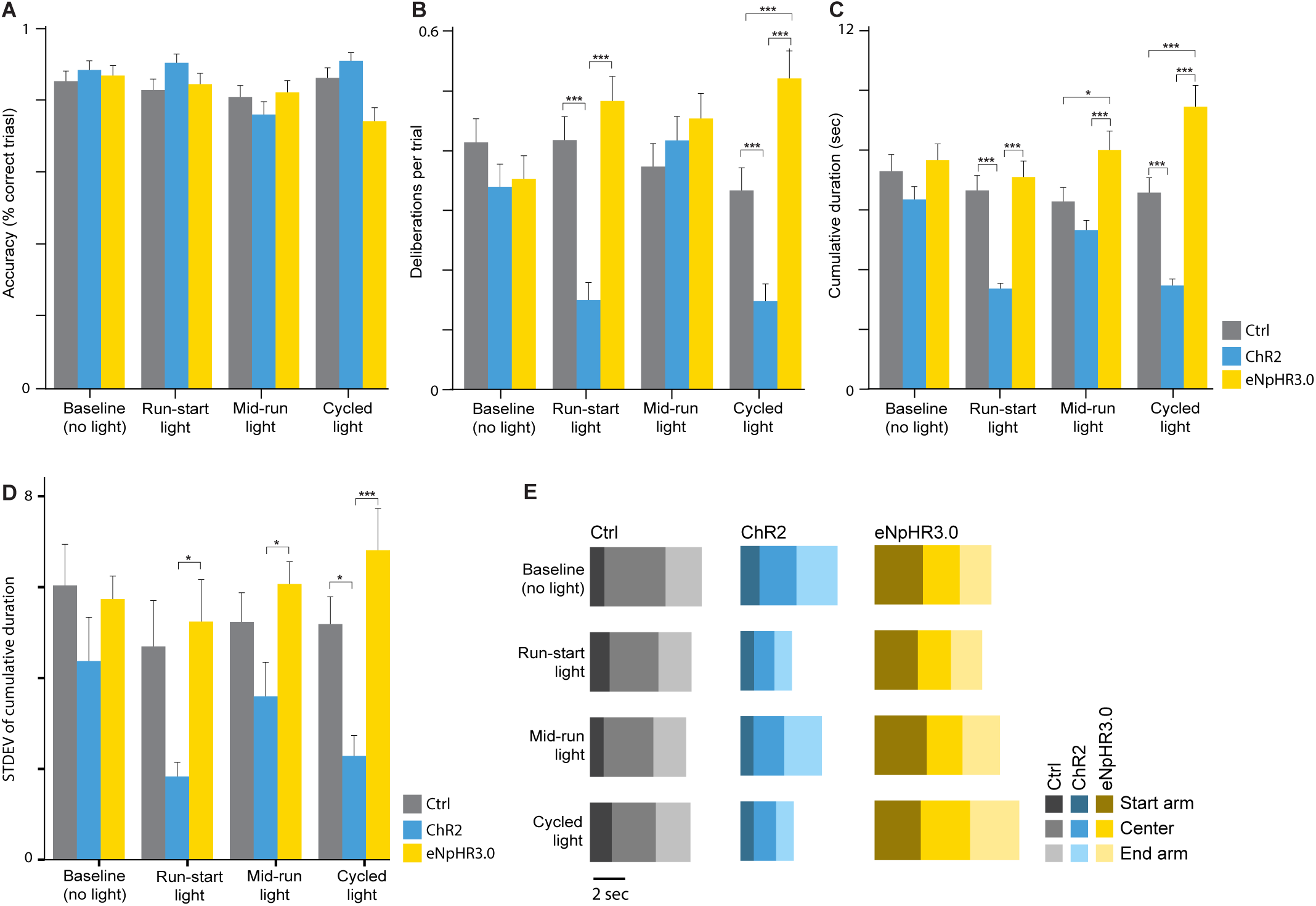
Changes in response maze behavior with DLS manipulations. **(A)** Percent correct running across the baseline and illumination test days. No main effects of Illumination (*F*_3,1017_=2.087, *P*=0.100) or Group (*F*_2,339_=1.073, *P*=0.343) nor interactions: Illumination/Group (*F*_6,1017_=1.641, *P*=0.133), Illumination/Trial (*F*_57,1017_=0.881, *P*=0.722), Illumination/Group/Trial (*F*_114,1017_=1.084, *P*=0.268). **(B)** Average deliberations per trial. Effect of Illumination (*F*_3,1014_=3.423, *P*=0.017), Group (*F*_2,338_=11.023, *P*<0.001), and Illumination/Group interaction (*F*_6,1014_=5.858, *P*<0.001). No interaction effects of Illumination/Trial (*F*_57,1014_=0.963, *P*=0.555) or Illumination/Group/Trial (*F*_114,1014_=1.030, *P*=0.402). Effects of Group during run-start (*F*_2,451_=22.148, *P*<0.001) and cycled (*F*_2,449_=25.975, *P*<0.001) illumination days. Posthocs showed the ChR2 group deliberated less than the control (*P*<0.001) and eNpHR3.0 (*P*<0.001) groups on the run-start illumination day. Behavior on cycled illumination days followed the same trends (ChR2/control: *P*<0.001; eNpHR3.0/control: *P*<0.001; ChR2/eNpHR3.0: *P*<0.001). **(C)** Cumulative duration of runs. Effect of Illumination (*F*_3,1014_=4.828, *P*=0.002) and Illumination/Group interaction (*F*_6,1014_=3.897, *P*<0.001). No interaction of Illumination/Trial (*F*_57,1014_=0.934, *P*=0.616) or Illumination/Group/Trial (*F*_114,1014_=1.081, *P*=0.273). Effect of Group (*F*_2,338_=17.992, *P*<0.001) on all three illumination days: run-start (*F*_2,451_=20.733, *P*<0.001), mid-run (*F*_2,447_=7.663, *P*<0.001), and cycled (*F*_2,449_=34.748, *P*<0.001). Post-hoc comparisons for run-start illumination showed that the ChR2 group was faster than controls (*P*<0.001) and eNpHR3.0 rats (*P*<0.001). Mid-run inhibition resulted in slower trials against controls (*P* =0.033) and ChR2 rats (*P*<0.001). Cycled illumination post-hoc comparisons showed bidirectional effects of stimulation and inhibition: ChR2/control (*P*<0.001), eNpHR3.0/control (*P*<0.001), and ChR2/eNpHR3.0 (*P*<0.001). **(D)** Standard deviation of the cumulative duration. Main effect of Illumination (*F*_3,63_=3.276, *P*=0.027) and Group (*F*_2,21_=8.157, *P*=0.002), but no Illumination/Group interaction (*F*_6,63_=1.425, *P*=0.219). Main effect of Group for run-start illumination (*F*_2,21_=5.088, *P*=0.016; ChR2/control: *P*=0.053; eNpHR3.0/control: *P*=0.883; ChR2/eNpHR3.0: *P*=0.019). Main effect of SD Group for mid-run illumination (*F*_2,21_=3.896, *P*=0.036; ChR2/control: *P*=0.189; eNpHR3.0/control: *P*=0.629; ChR2/eNpHR3.0: *P*=0.031). Main effect of Group for cycled illumination (*F*_2,21_=11.183, *P*<0.001; ChR2/control: *P*=0.018; eNpHR3.0/control: *P*=0.239; ChR2/eNpHR3.0: *P*<0.001). **(E)** Duration of running in maze segments (start arm, center, end arm). Rows denote illumination day. Columns denote treatment group. Percent duration differences for each maze segment for ChR2 vs. controls: 1) Baseline = 14% slower in the start arm, 25% faster in the center, and 7% slower at the end arm; 2) Run-start light = 18% faster in the start arm, 41% faster in the center, and 31% faster at the end arm; 3) Mid-run light = 2% slower in the start arm, 23% faster in the center, and 7% slower at the end arm; 4) Cycled light = 24% faster in the start arm, 33% faster in the center, and 33% faster at the end arm. Percent duration differences for each maze segment for eNpHR3.0 vs. controls: 1) Baseline = 54% slower in the start arm (reflecting a group predisposition), 24% faster in the center, and 6% faster at the end arm; 2) Run-start light = 38% slower in the start arm, 19% faster in the center, and 3% faster in the end arm than control; 3) Mid-run light = 58% slower in the start arm, 18% faster in the center, and 6% slower in the end arm than control; 4) Cycled light = 36% slower in the start arm, 5% slower in the center, and 18% slower in the end arm. Statistics for **E** are in Supplementary Table 1. For all figure panels, bars and errors show mean ± SEM; asterisks denote significant post-hoc comparisons (∗*P*<0.05, ∗∗*P*<0.01, ∗∗∗*P*<0.001); lack of asterisks denotes lack of significance.

The accuracy of the runs was not affected by DLS run-start stimulation (Fig. 2A), suggesting that the use of task rules to inform behavior was unaffected. However, accuracy was reduced when DLS was inhibited on the cycled schedule, suggesting some sensitivity of accuracy to DLS disruption. Regardless, the result shows that the vigor of runs on a DLS-dependent response task are powerfully modulated by DLS activity at the initiation point of behavior, and this vigor-promoting function is dissociable from the execution of behavior based on a turn-rule or stimulus-response association, at least as they are indexed by performance accuracy.

We next examined how the change in run vigor produced by run-start manipulations related to habit strength by using a reward devaluation procedure. This procedure included tests of how changes in the value of the expected outcome (unrewarded extinction conditions) and received outcome (rewarded reacquisition session) were used in behavior (Fig. 3). A more (or less) habitual run would exhibit independence from (or sensitivity to) changes in the outcome value. We first established a baseline by giving animals a probe test under unrewarded conditions with DLS light delivery at run-start, followed by a normal rewarded reminder session (Fig. 3A). Then, free intake of the maze reward in home-cages was paired with nauseogenic lithium chloride until an aversion to the reward developed and it was rejected by the rats. Rats were then returned to the task and exposed to a post-devaluation unrewarded probe test followed by a post-devaluation rewarded test; both with DLS light being delivered at run-start.

**Figure 3.**
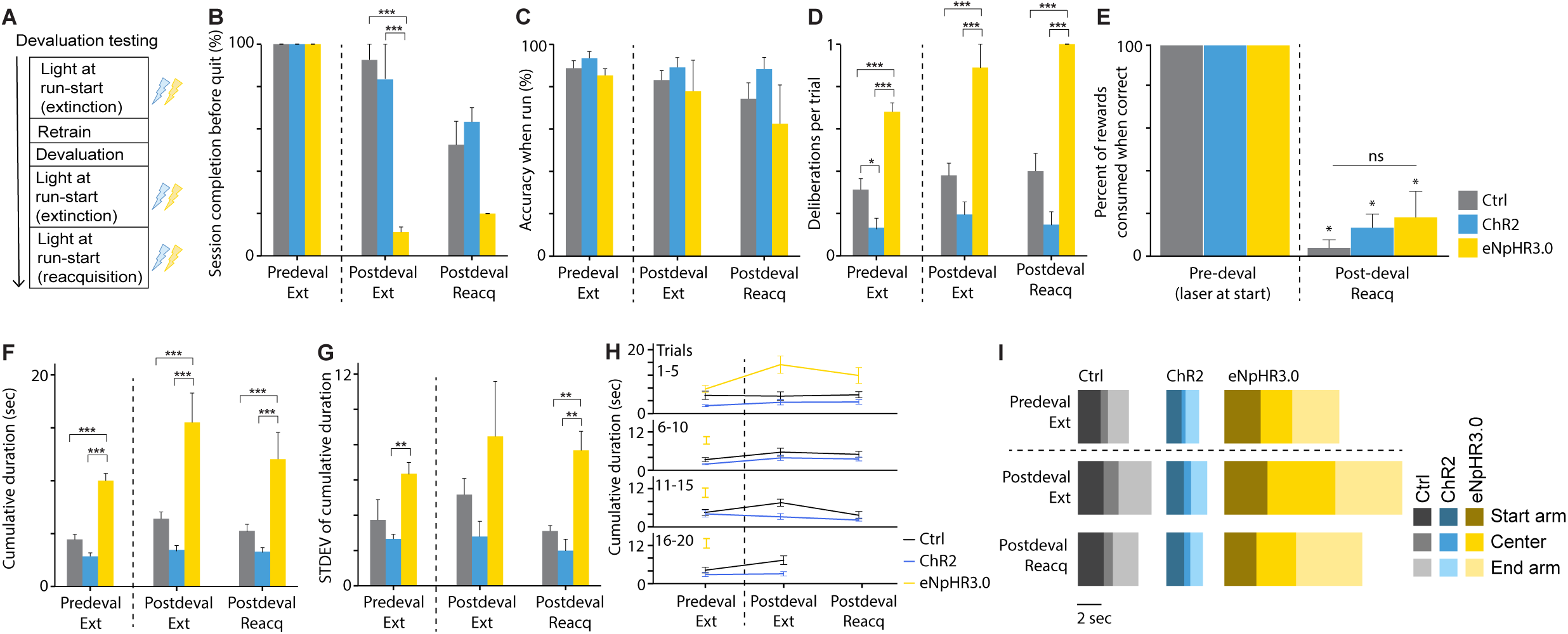
Response maze with reward devaluation. **(A)** Devaluation testing timeline. **(B)** Percent trials completed. Between pre- and post-devaluation extinction days, main effect of Illumination (*P*<_1,24_=64.444, *P*<0.001) and Group (*F*_2,24_=35.008, *P*<0.001), and Illumination/Group interaction (*F*_2,24_=35.008, *P*<0.001). Posthocs showed this to be due to eNpHR3.0 rats quitting more after devaluation (chR2/eNpHR3.0 *P*<0.001; control/eNpHR3.0 *P*<0.001, ChR2/Control *P*=0.735). The same trend, though non-significant as all rats quit more, occurred during reacquisition (*F*_1,9_=0.117, *P*=0.077). **(C)** Accuracy when rats ran. There were no significant differences between extinction days for Illumination (*F*_1,123_=3.187, *P*=0.077), Group (*F*_2,123_=0.819, *P*=0.443) or Illumination/Group (*F*_2,123_=0.562, *P*=0.572) or in the post-devaluation rewarded day for Group (*F*_2,75_=1.309, *P*=0.276). **(D)** Deliberations per trial. No main effect across pre-devaluation extinction versus post-devaluation extinction for Illumination day (*F*_1,94_=2.436, *P*=0.122), nor interactions: Illumination/Group (*F*_2,94_=0.145, *P*=0.865); Illumination/Trial (*F*_19,94_=0.521, *P*=0.947); Illumination/Group/Trial (*F*_27,94_= 0.716, *P*=0.838). Main effect of Group on all three illumination days: pre-devaluation extinction (*F*_2,203_=32.478, *P*<0.001; control/ChR2 *P*=0.050, control/eNpHR3.0 *P*<0.001, ChR2/eNpHR3.0 *P*<0.001); post-devaluation extinction: (*F*_2,94_=23.185, *P*<0.001; control/eNpHR3.0 *P*<0.001; ChR2/eNpHR3.0 *P*<0.001); post-devaluation reacquisition: (*F*_2,52_=21.198, *P*<0.001; control/eNpHR3.0 *P*<0.001; ChR2/eNpHR3.0 *P*<0.001). **(E)** Percent of rewards consumed on task prior to quitting a run, which were reduced across groups and trials from pre- to post-devaluation with illumination at run-start. Effects of Illumination day (*F*_1,356_=2776.301, *P*<0.001) and interactions: Illumination/Trial (*F*_14,356_=7.981, *P*<0.001), Group/Illumination (*F*_2,356_=7.981, *P*<0.001), Illumination/Group/Trial (*F*_25,356_=4.434, *P*<0.001). No main effect of Group on pre-devaluation day (*F*_2,262_=0) or reacquisition (*F*_2,94_=2.177, *P*=0.119). **F)** Cumulative duration of runs. Main effect of Illumination day (*F*_1,98_=74.492, *P*<0.001) and Illumination/Group interaction (*F*_2,98_=28.436, *P*<0.001). No interaction of Illumination/Trial (*F*_19,98_=0.795, *P*=0.708) nor Illumination/Group/Trial (*F*_28,98_=0.955, *P*=0.537). Main effect of Group on all three illumination days: Pre-devaluation extinction (*F*_2,202_= 41.428, *P*<0.001; eNpHR3.0/control *P*<0.001; eNpHR3.0/ChR2 *P*<0.001; control/ChR2 *P*=0.237); post-devaluation extinction (*F*_2,99_=69.088, *P*<0.001; eNpHR3.0/control *P*<0.001; eNpHR3.0/ChR2 *P*<0.001; control/ChR2 *P*=0.065); post-devaluation reacquisition (*F*_2,55_=23.010, *P*<0.001; eNpHR3.0/control *P*<0.001; eNpHR3.0/ChR2 *P*<0.001; control/ChR2 *P*=0.538). **(G)** Standard deviation of the cumulative duration. Comparing extinction days, no effect of Illumination day (*F*_1,24_=1.388, *P*=0.254), an effect of Group (*F*_2,24_=6.933, *P*=0.006), and no Group/Illumination interaction (*F*_2,24_=0.290, *P*=0.751). Group effect driven by greater variability in eNpHR3.0 than ChR2 groups (eNpHR3.0/ChR2 *P*=0.009; eNpHR3.0/control *P*=0.098; ChR2/control *P*=0.407). Greater duration variability in eNpHR3.0 rats continued in post-devaluation reacquisition (*F*_2,9_=21.166, *P*=0.002; control/eNpHR3.0 *P*=0.004, ChR2/eNpHR3.0 *P*=0.002; control/ChR2 *P*=0.369). **(H)** Cumulative duration per trial block. Trial did not interact with illumination effects, as above. **(I)** Duration of running in maze segments (start arm, center, end arm). Rows denote illumination day. Columns denote treatment group. Dotted line represents reward devaluation. Percent duration differences for each maze segment ChR2 vs. controls: 1) Pre-devaluation extinction = 20% faster in the start arm, 30% faster in the center, and 21% faster at the end arm. 2) Post-devaluation extinction = 19% faster in the start arm, 36% faster in the center, and 34% faster in the end arm. 3) Post-devaluation reacquisition = 18% faster in the start arm, 21% faster in the center, and 34% faster in the end arm. Percent duration differences for each maze segment eNpHR3.0 vs. Control: 1) Pre-devaluation extinction = 23% slower in the start arm, 62% slower in the center, and 38% slower in the end arm; 2) Post-devaluation extinction = 46% slower in the start arm, 70% slower in the center, and 49% slower in the end arm; 3) Post-devaluation reacquisition = 21% slower in the start arm, 79% slower in the center, and 58% slower in the end arm. Statistics are in Supplementary Table 2. For all panels, asterisks denote significant posthoc comparisons (^∗^*P*<0.05, ^∗∗^*P*<0.01, ^∗∗^*P*<0.001, ‘ns’ not significantly different).

Control animals behaved habitually, running to the devalued reward location in the unrewarded probe test at a level of accuracy that was comparable to the pre-devaluation probe test (Fig. 3C). This produced a potential ceiling for increasing habit strength. In the rewarded test, control animals began avoiding the devalued arm by neglecting to perform the task (Fig. 3B), and when they did run the task they rejected nearly all of the rewards (Fig. 3E). This result confirmed that the devaluation memory generalized appropriately to the task. On each test day (pre- and post-devaluation probe; post-devaluation reacquisition), DLS stimulation at run-start produced a similar pattern of vigor enhancement by way of reducing run duration (Fig. 3F), run duration variance (Fig. 3G), and deliberations (Fig. 3D), which was not limited to behavior at the initial maze segment (Fig. 3H, 3I; Supplementary Table 2). At the same time, the level of neglecting to run the task, run accuracy when they did run, and level of reward rejection were all similar to controls. In contrast, animals with DLS inhibition at run-start exhibited evidence of vastly reduced run vigor and loss of habit. On each test day, including the pre-devaluation probe, their runs (when they did run) were far slower and more variant, and deliberations occurred on nearly every trial. Moreover, these animals essentially quit the task after reward devaluation; they neglected to execute runs almost immediately in the task after the reward was devalued in the unrewarded probe session. Their rejection of rewards when they did run was similar to controls. Again, the effect on running behavior was not simply restricted to the run-start period in which the manipulation was given (Supplementary Table 2).

This set of results reveals several important features of DLS function at run-start: (1) it modulates performance vigor bidirectionally, but vigor reduction via DLS inhibition is most evident when the outcome of the runs is reduced (unrewarded or devalued reward conditions); (2) its activation increases the vigor of outcome-insensitive (habitual) actions, and its inhibition decreases both vigor and outcome sensitivity; (3) it does not affect the valuation of the reward itself as indicated by reward rejection levels. In other words, greater DLS signaling at the initiation of behavior improves vigor in conditions when outcomes are stable or when they change, while DLS inhibition reduces vigor and in turn increases purposefulness (i.e., outcome sensitivity) preferentially when outcome values change. This latter result is reminiscent of habit loss (i.e., outcome sensitivity gain) with prior DLS loss-of-function work^1,2^, and potentially pinpoints this result to compromised DLS signaling at run-start specifically. Importantly, the post-devaluation effects of eNpHR3.0 show that the lack of change in run accuracy before reward devaluation in the same animals is unlikely to have been due to eNpHR3.0 ineffectiveness.

These results raised the question of whether the improvement in vigor that was observed with run-start, but not mid-run, DLS stimulation was because that was the point in time at which a full run routine was selected and set in motion. To test this, we developed a ‘beacon’ plus-maze task, derived from the DLS-dependent win-stay task^34-37^, where a new cohort of rats learned to navigate towards a visual cue for reward (Fig. 4A). The cue could only be ascertained at the middle choice point and its location was varied across trial blocks; it could not be reliably associated with a turn direction or with extra-maze cues. Thus, this task required an action to be selected at the start (i.e., run to the middle) and again at the detection of the cue mid-behavior (i.e., run to cue for reward). We focused on comparing ChR2-mediated DLS stimulation to controls (Fig. 4C). An identical training (Fig. 4D) and manipulation protocol as above was used (Fig. 4B).

**Figure 4.**
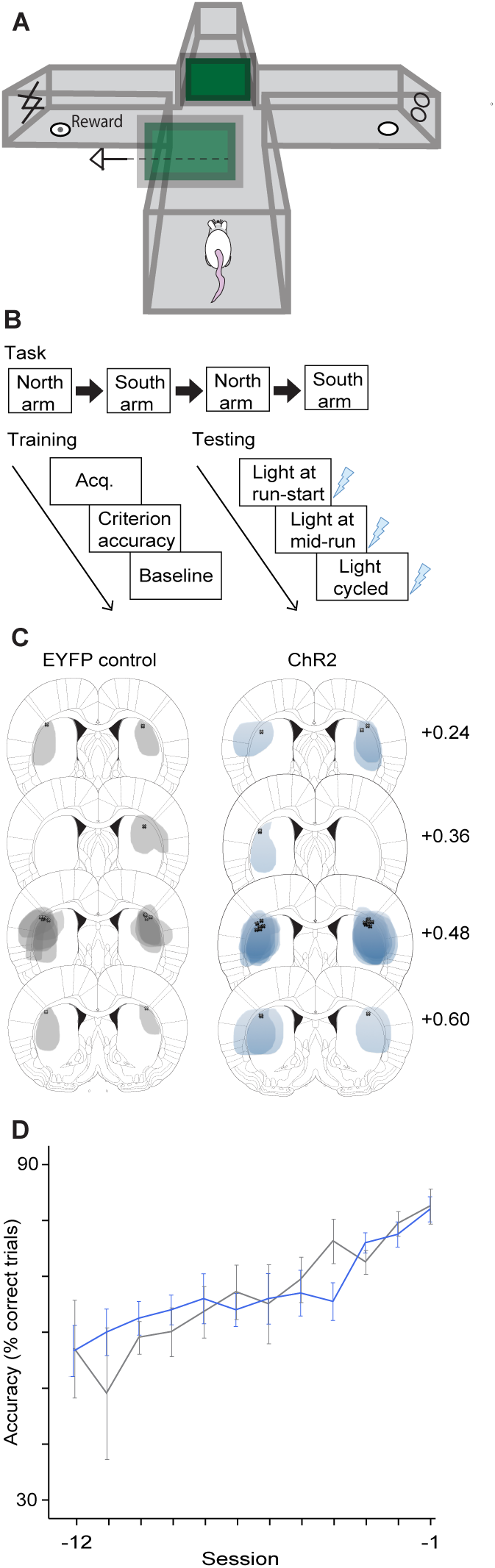
Beacon task. **(A)** Maze setup and laser manipulation protocol. Task protocol (same as Fig. 1A). **(B)** Expression mapping of control and ChR2 groups, produced as in Fig. 1C. **(C)** Run accuracy in sessions leading up to the final day of criterion performance, separated by ChR2 (blue) and control (gray) groups. No main effect of Group (*F*_1,258_=1.018, *P*=0.314) or Group/Day interaction (*F*_28,158_= 0.914, *P*=0.574), but main effect of Day (*F*_21,253_=5.514, *P*<0.001).

Increased vigor was replicated in our beacon task condition with DLS stimulation at run-start. This manipulation reduced run duration (Fig. 5C) and deliberations (Fig. 5B), without affecting task accuracy (Fig. 5A). Remarkably, in contrast to the response task, mid-run DLS stimulation was suddenly also effective at improving overall vigor on this task. Enhancing DLS activity at this point in the run, which was when the cue was located and approached, also increased vigor at comparable levels to the increase caused by run-start stimulation. Cycled stimulation was essentially identical as well. No clear restrictions of the effect to particular maze segments were observed (Fig. 5E; Supplementary Table 3). This result points to a phasic DLS signaling role in performance vigor at the time major action choices are made.

**Figure 5.**
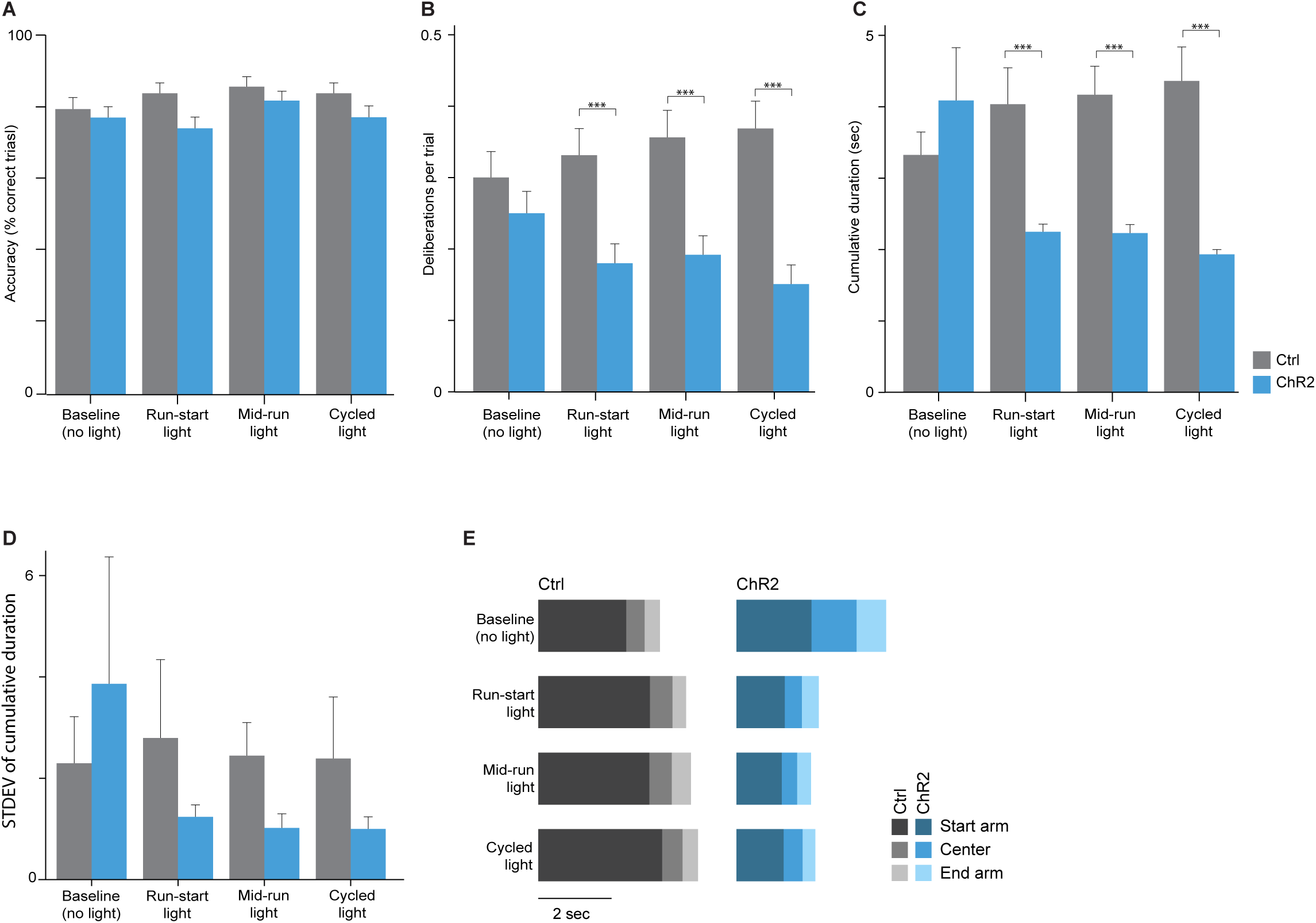
Changes in beacon maze running behavior. **(A)** Percent correct trials on beacon task. No main effects for Illumination (*F*_3,954_=1.531, *P*=0.205) nor interactions: Illumination/Group (*F*_3,954_=0.703, *P*=0.550), Illumination/Trial (*F*_57,954_=0.791, *P*=0.868), Illumination/Group/Trial (*F*_57,954_ =1.107, *P*=0.277). Main effect of Virus with illumination at run-start only (*F*_1,320_=4.916, *P*=0.027). **(B)** Deliberations per trial. No main effect of Illumination (*F*_3,954_=0.229, *P*=0.876) nor interactions of Illumination/Trial (*F*_57,954_=1.282, *P*=0.082) or Illumination/Group/Trial (*F*_57,954_=0.823, *P*=0.822). There was an Illumination/Group interaction (*F*_3,954_=2.622, *P*=0.050). No effect of Group on the baseline day (*F*_1,320_=1.111, *P*=0.293), but significant differences during illumination at run-start (*F*_1,320_=11.381, *P*<0.001), mid-run (*F*_1,319_=18.376, *P*<0.001), and cycled (*F*_1,319_=17.367, *P*<0.001). **(C)** Cumulative duration of runs. No main effect of Illumination (*F*_3,954_=1.190, *P*=0.312), nor interactions of Illumination/Trial (*F*_57,954_=1.102, *P*=0.285) or Illumination/Group/Trial (*F*_57,954_=0.565, *P*=0.996). There was an Illumination/Group interaction (*F*_3,954_=7.727, *P*<0.001). No effect of Group on the baseline day (*F*_1,320_=0.739, *P*=0.390), but significant differences during run-start (*F*_1,320_=13.729, *P*<0.001), mid-run (*F*_1,319_=30.453, *P*<0.001), and cycled illumination days (*F*_1,319_=23.895, *P*<0.001). **(D)** Standard deviation of the cumulative duration. Not significant for Illumination (*F*_3,48_=0.856, *P*=0.470), Group (*F*_1,16_=0.311, *P*=0.585), or Illumination/Group (*F*_1,16_=1.349, *P*=0.262). **(E)** Duration of running in maze segments (start arm, center, end arm). Rows denote illumination day. Columns denote treatment group. Percent duration differences for each maze segment ChR2 vs. Control: 1) Baseline: 8% faster in the start arm, 32% slower in the center, and 42% slower in the end arm; 2) Run-start light: 40% faster in the start arm, 10% slower in the center, and 14% faster in the end arm; 3) Mid-run light: 42% faster in the start arm, 16% faster in the center, and 18% faster in the end arm; 4) Cycled light: 45% faster in the start arm, 10% faster in the center, and 4% faster in the end arm. Statistics in Supplementary Table 3. For all panels, asterisks denote significant posthoc comparisons (^∗^*P*<0.05, ^∗∗^*P*<0.01, ^∗∗^*P*<0.001).

The reward devaluation protocol then followed (Fig. 6A). Control animals ran this task habitually by electing to run nearly all trials (Fig. 6B, 6C), and by running to the devalued reward locations in the postdevaluation probe test. Intriguingly, they also continued to run accurately in the rewarded reacquisition test that followed, despite mostly rejecting the rewards (Fig. 6E). This level of persistence could reflect an incentive attraction to the maze cue itself rather than the paired outcome^38,39^. However, there were still improvement effects on the vigor of runs from DLS stimulation. In the post-devaluation unrewarded probe test, run-start stimulation of the DLS reduced run duration (Fig. 6F) and deliberations (Fig. 6D) compared to controls. One exception was lack of vigor change during the post-devaluation reacquisition session, which could be attributed to the realization of the paired aversion, driven by later session trials 16-20 (Fig. 6H). Behavior changes were again not limited to particular maze running segments (Fig. 6I; Supplementary Table 4). The consumption/rejection of the rewards themselves was unaffected by DLS stimulations (Fig. 6E).

**Figure 6.**
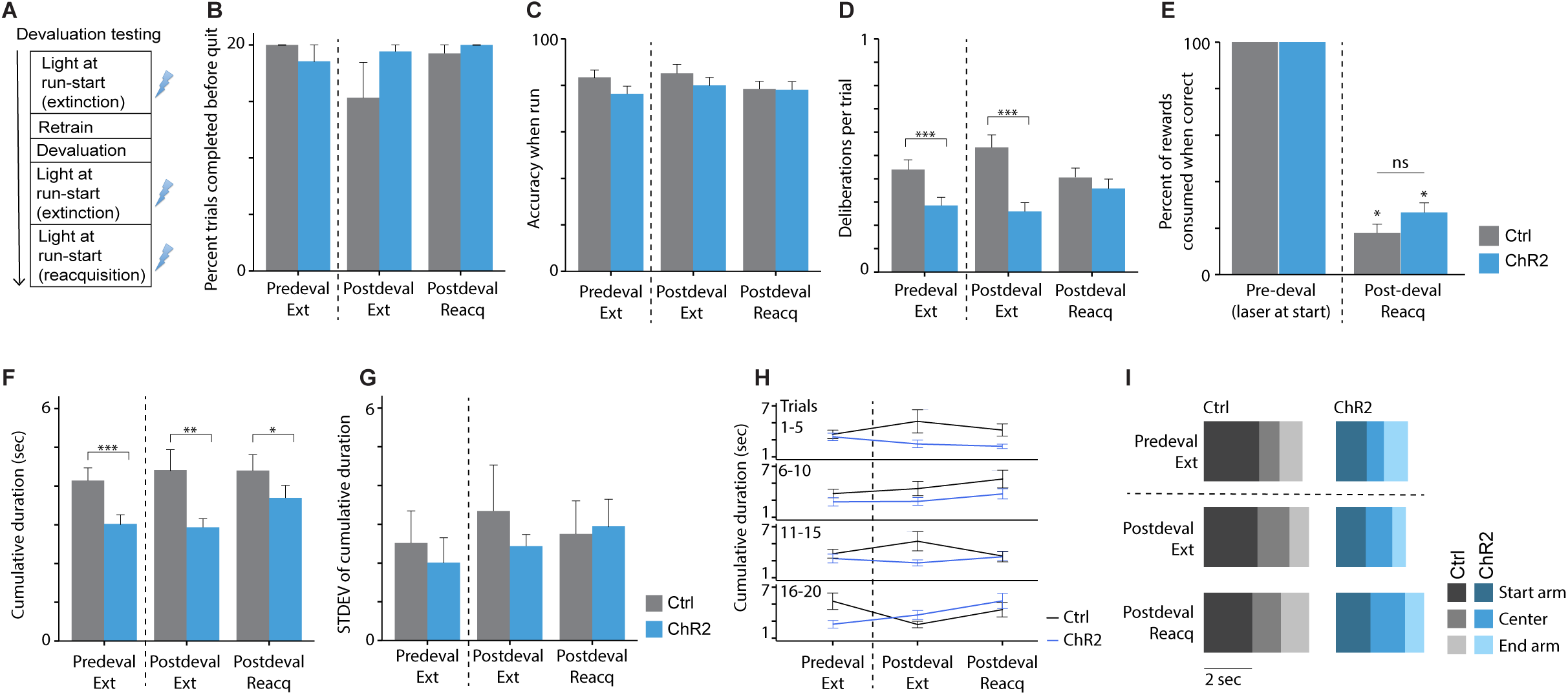
Beacon task with reward devaluation. **(A)** Devaluation testing timeline. **(B)** Percent trials completed, which remained high and stable. Between pre- and post-devaluation extinction days, no main effect of Illumination day (*F*_1,30_=0.108, *P*=0.746) and Group (*F*_1,30_=0.460, *P*=0.504), nor Illumination/Group interaction (*F*_1,30_=0.034, *P*=0.854). No effect of Group during reacquisition (*F*_1,16_=0.450, *P*=0.513). **(C)** Performance accuracy when rats ran, which was also stable. Between pre- and post-devaluation extinction days, no main effect of Illumination (*F*_1,214_=2.135, *P*=0.145), Group (*F*_1,214_=0.464, *P*=0.497) or Illumination/Group interaction (*F*_1,214_=0.169, *P*=0.682). On the reacquisition day no effect of Group (*F*_1,270_=0.035, *P*=0.852). **(D)** Deliberations per trial. No main effect of Illumination (*F*_1,214_=0.314, *P*=0.576) or interactions: Illumination/Group (*F*_1,214_=0.183, *P*=0.669), Illumination/Trial (*F*_19,214_=0.672, *P*=0.844), Illumination/Group/Trial (*F*_19,214_=0.931, *P*=0.545). Main effects of Group in both extinction illumination devaluation day: pre-devaluation extinction (*F*_1,297_=11.142, *P*<0.001); post-devaluation extinction (*F*_1,215_=16.901, *P*<0.001). No Group effect in post-devaluation reacquisition (*F*_1,270_=0.785, *P*=0.376). **(E)** Percent of rewards consumed on task prior to quitting a run, which were reduced equivalently across groups from pre- to post-devaluation with illumination at run-start. Effects of Illumination day (*F*_1,627_=1726.655, *P*<0.001) and interactions: Illumination/Trial (*F*_19,627_=2.875, *P*<0.001), Illumination/Group/Trial (*F*_19,627_=1.829, *P*=0.017). No Group/Illumination interaction (*F*_1,356_=2.667, *P*=0.103). No main effect of Group on pre-devaluation day (*F*_1,337_= 0) or reacquisition (*F*_1,290_=1.944, *P*=0.164). **(F)** Cumulative duration of runs. No main effect for Illumination (*F*_1,214_=0.514, *P*=0.474), or interactions: Illumination/Trial (*F*_19,214_=1.571, *P*=0.065), Illumination/Group/Trial (*F*_19,214_=0.990, *P*=0.475). There was an Illumination/Group interaction (*F*_1,214_=5.644, *P*=0.018). Main effects of Group on in all three illumination days: pre-devaluation extinction (*F*_1,297_=15.331, *P*<0.001); post-devaluation extinction (*F*_1,215_=7.671, *P*=0.006); post-devaluation reacquisition (*F*_1,270_=4.455, *P*=0.036). **(G)** Standard deviation of the cumulative duration. No main effect of Illumination (*F*_1,11_=0.674, *P*=0.429), Group (*F*_1,11_=0.475, *P*=0.505) or Illumination/Group interaction (*F*_1,11_=3.283, *P*=0.097). Post-deval reacquisition no effect of Group (*F*_1,14_=0.271, *P*=0.611). **(H)** Cumulative duration per block. No significant trial interactions as above. **(I)** Duration of running in maze segments (start arm, center, end arm). Rows denote illumination day. Columns denote treatment group. Dotted line represents reward devaluation. Though there were no trial interactions, there was trend towards faster runs in ChR2 rats during early reacquisition blocks. Percent duration differences for each maze segment ChR2 vs. Control: 1) Pre-devaluation extinction: 60% faster in the start arm, 5% faster in the center, and 11% faster in the end arm; 2) Post-devaluation extinction: 45% faster in the start arm, 3% faster in the center, and 11% faster in the end arm; 3) Post-devaluation reacquisition: 38% faster in the start arm, 13% faster in the center, and 9% faster in the end arm. Statistics in Supplementary Table 4. For all panels, asterisks denote significant posthoc comparisons (^∗^*P*<0.05, ^∗∗^*P*<0.01, ^∗∗^*P*<0.001, ‘ns’ not significantly different).

## Conclusions

In all, these findings show that phasic DLS signaling promotes the vigor of response-based and cue-based behaviors at the time they are selected or initiated, which is dissociable from possible DLS roles in the use of reinforcement rules to accurately perform these tasks. Moreover, phasic DLS activity appears to play a key role in promoting vigorous execution of habits, and loss of this DLS activity leads to a behavior that is reduced in vigor and is more goal-directed. In contrast, whether a learned behavior occurs as a habit or as a purposeful goal-directed action is thought to be be dictated by the level of participation amongst competing cortico-basal ganglia neural systems that are thought to promote one or the other strategy^6,7,40,41^. Our results help show that the vigor of a behavior, and how outcome sensitive it is can be modulated by increased levels of phasic DLS activity itself^42-44^. We therefore propose a habit-by-way-of-vigor function for the DLS and the broader basal ganglia that departs from traditional rule/association learning frameworks for understanding habits. We favor this over another possible explanation of our results, namely that DLS activity suppresses outcome processing giving vigor as a default, due to the much closer correspondence of DLS firing activity to run speed and fluidity rather than to outcome sensitivity^18^. One possible way in which changes in vigor could directly alter habit strength is to gate the ability of outcome evaluation processes to affect behavior by changing the temporal window that would allow or restrict them from occurring; when vigor is high over a succession of performance runs, greater outcome insensitivity will result. Finally, our results suggest the importance of basal ganglia areas in controlling effortful behavior for reward in a manner that could be similar to what has been described of mesolimbic systems^45-49^

## Figure legends

**Supplementary Figure 1**. Histograms (0.02 sec bins) of all single recorded units that were responsive to blue light. Firing rate (Hz) is shown 0.5 sec before, during, and after light delivery. Left column section, units recorded under anesthesia (n = 9). Right column section, units recorded during free behavior (n = 3). Rows within column show the same unit recorded when light was delivered 0.5 sec every 1 min (left subcolumn) and when light was delivered in a 0.5 sec on/off cycle (right subcolumn). Gaps denote units that were not stable or were not detected during that light delivery protocol. Although not all units responded to light immediately, they nonetheless exhibited response excitation as confirmation of ChR2-mediated stimulation of the DLS. Units with activity inhibited by light were not observed.

## Materials and Methods

### Subjects and Surgery

Male Long Evans rats (n = 45) were individually housed in a dedicated animal vivarium under direct care procedures approved by the Dartmouth College Institutional Animal Care and Use Committee. All rats were obtained at 250-400 g and then maintained on a 85% post-surgical weight for the training and testing duration. Rats were housed on a reverse light-dark cycle, with response task experiments conducted in the dark and beacon task experiments conducted in the light. Surgical procedures were performed using aseptic techniques under isoflurane anesthesia: intracranial injection of viral vectors, intracranial implantation of fiber optic guides, and implantation of tetrode-carrying headstages (VersaDrive-8 Optical, Neuralynx freely moving and anesthetized recordings: n = 3). Rats received bilateral injections (0.3 μL volume) into DLS using a microinfusion pump and 33-gauge syringe of a single viral construct: channelrhodopsin-2 (response task: n = 8, beacon task: n = 10; AAV5-hSYN-ChR2-EYFP); eNpHR3.0-halorhodopsin (response task: n = 8, AAV5-hSYN-eNpHR3.0-EYFP); sham control (response task: n = 8, beacon task: n = 8, AAV5-hSYN-EYFP). Bilateral DLS injection and fiber implant coordinates were, in mm: AP +0.5, ML −/+ 4.0, DV injection −4.3 from skull, with fiber implants (200 μm, ThorLabs or in-house) terminating at DV −3.8 mm. Fiber and tetrode implants were permanently affixed to the skull using dental cement and skull screws. Rats in both tasks received fiber implants and viral vector infusions in a single surgery, and rats used in the *in vivo* electrophysiology experiments required two surgeries. All rats had at least one-week post-surgery recovery time.

### Experimental Procedures

Male Long Evans rats (total n = 42) were exposed to a custom-built plus maze on two separate motor action sequence tasks for reward. First, the response task rats (n = 24) learned to turn the same direction (i.e., always turn right at a choice point) towards a baited end arm for a single pellet reward, regardless of starting maze location. In total, 20 turns (e.g., all right) would be rewarded in a daily session with turn direction counterbalanced across animals. Rats in this task had the opportunity to correct themselves if they did not enter the correct arm on the first try, with a total of 30 sec to complete a trial. A circuitous (i.e., multiple arm entries) but ultimately accurate run was still recorded as an incorrect trial when calculating overall accuracy due to the lack of a clear response strategy in the run. Trials exceeding 30 sec were counted as ‘misses’ and excluded from analysis. In the event of a miss, the session would continue until 20 trials were complete. After devaluation, on occasion a rat would refuse to run several trial opportunities in a row (a tendency of the eNpHR3.0 group), in which case the session was concluded. To account for this, data were assessed for trials leading up to the first failed trial opportunity. Training progressed over daily sessions until acquisition of performance accuracy reached (≥75% for three days). DLS illumination testing days followed. Habitual runs were tested operationally using a pre-extinction day, followed by a reward devaluation procedure (e.g., 3 separate pairings of LiCl injections following 20-30 min free reward period in home-cage in a separate context), a post-devaluation extinction probe day, and a post-devaluation rewarded (reacquisition) day. All behavior testing was performed and recorded in the dark under a red light to minimize the use of visual cues. Run durations per maze segment, and overall, were calculated. Deliberations were counted as either occurring or not (max 1/trial) using a XY-frame of reference for the nose-point at the center region of the maze. Deliberations included a clear turn of the head down one maze arm and then a return of the head to be forward facing or toward the alternate arm. For analysis of percent trials completed within a session before quitting (an effect of reward devaluation), the first trial a rat failed to complete within 30 sec was used (e.g., if it was trial 10 out of 20, then the measure was 50%).

In the beacon task, rats (n = 18) learned to follow an end arm cue (two vertical XX’s or OO’s, Fig. 4A) that was paired with a reward pellet. Starting location and reward-paired cues were arranged in such a way to avoid response actions (i.e., always turn left at choice point). In total, daily sessions had 20 trials with an equal number of right and left turns (e.g., 5 right, 5 left, 5 left, 5 right). The beacon task was run in the light so that the task cues could inform performance. Behavioral monitoring and training/testing protocols otherwise followed identically from the response task.

### Plus Maze Apparatus Setup and Training

A Plexiglas plus maze was used for both the response and beacon maze experiments. Four total maze arms were identical in length (35.56 cm), width (17.78 cm), and height (33.02 cm), separated by a square center region (diameter: 25.4 cm) (Fig. 1A, 4A). For each trial, a removable insert blocked off the arm opposite to the starting arm location. Rats had two directions (i.e., turn left or right) to choose from at a middle choice point. End arms had identical reward dishes, with one baited with a reward pellet and the other empty.

Prior to training, rats were exposed to the plus maze for a 30 min habituation session, proceeded by home-cage free exposure of reward pellets overnight. Daily sessions of 20 trials followed: rats would await a trial in one of two starting arm locations (e.g., north arm or south arm, counterbalanced) that was blocked off by a removable insert separating the starting arm from the runway; once the insert was removed (counterbalanced as left or right so as not to bias the rat to turn in one direction), signaled the rat a trial had begun. The rat then traversed the starting arm runway to the maze center decision point, having an equal choice (50%) of turning left or right. However, only one end arm was baited with reward and this assignment remained consistent per rat (i.e., response: turn direction-reward; beacon: cue-reward).

The difference between the two tasks can be seen by the action required for reward. For the response task, the reward location was consistent across all trials independent of starting arm, requiring the same action (e.g., right turn) for reward. 20 trials were conducted on a daily session with a maximum of 20 rewards. Each session was a block design. 5 trials would start from the north arm and require a rat to run and turn right for reward; the animal would then be manually placed in the south arm, requiring the rat to turn right again for reward for 5 trials. The rat would be placed back in the north arm again for 5 trials, ending with an additional 5 trials from the south arm. After each trial (i.e., either pellet was consumed as a hit or corrected hit, or a miss (no consumption): 30 seconds max to find reward), the rat would be placed back into the starting arm. There was a ~1 min interval between trials. Therefore, a perfect run would be 20 correct right turns from either a north or south arm starting location. The assigned turn direction yielding reward was pseudorandomly assigned across animals and groups.

For the beacon task, one of two removable cues (two vertical ‘XX’s or ‘OO’s) was affixed to one of two end maze arms that could be switched. One cue was paired with reward at one end arm, with the other cue paired with an empty dish in the opposite arm that also stayed consistent across trials. Neither of the cues could be detected until the center region was reached due to the tall opaque maze walls that were covered with a green removable tarp that could be affixed directly to the maze and easily removed. As in the response task, rats began either in the north or south arm. Removal of the gate alerted the rat a trial had begun. The rat then traversed the runway to the center, where the decision (turn left or right) was made based on location of the reward-paired cue. Incorrect turns to the unpaired cue resulted in no reward. Unlike the response task, rats that chose the incorrect, unpaired-cue arm were not allowed to correct themselves. For instance, a Plexiglas insert was used to block off the rat from entering the correct arm if they chose the unpaired-cue arm first, resulting in manually placing the rat back to the starting arm location. In this way, rats learned to locate the cue predicting reward rather than just guessing and exploring both end arms and not making the association that one of two cues was rewarded and the other resulted in nothing. In contrast to the response task, multiple major actions were required for this task: at the start (run to middle) and again at the center (run to reward-paired cue). In daily sessions, rats began in the same north or south arm locations. Sessions consisted of 20 trials per day with a maximum of 20 rewards, with a ~1 min interval between trials. A block design was also used (5-north, 5-south, 5-north, 5-south). Either the “XX” or “OO” cue would signify reward and stay in the same west or east arm for the first 10 trials and then shift to the alternate arm for the remaining 10 trials. All parameters were assigned pseudorandomly across rats. Training on this task also continued until the accuracy criterion, followed by illumination test days, and then the devaluation testing protocol exactly as administered in the response task.

### Optogenetic Manipulations

Optogenetic constructs were introduced into bilateral DLS through injections of AAV5-hSYN-EYFP (Control), AAV5-hSYN-ChR2-EYFP (Channelrhodopsin-2), or AAV5-hSYN-eNpHR3.0-EYFP (Halorhodopsin). Patch cables with 0.2 μm fiber core and 0.39 NA were used for all testing (Thorlabs, Newton, NJ; Doric Lenses, Quebec City, Canada). On DLS manipulation days, either 473 nm or 593.5 nm light from DPSS lasers (Shanghai Laser & Optics Century Co., Ltd.) was delivered through a fiber patch cord connected to an integrated rotary joint beam splitter (Thorlabs, Newton, NJ; Doric Lenses, Quebec City, Canada) that allowed for two fiber patch cords to be connected directly to surgically implanted optic fiber cannula (in-house fiber implants, 200 μm, ThorLabs) through ceramic sleeves terminating bilaterally into DLS. Laser delivery was gated either by Ethovision XT software (Noldus Inc.) or through a Master 9 pulse generator (A.M.P.I.). Power output was measured to be between 3-5 mW at the level of each patch cable ferrule connector per hemisphere prior to and after each test day. Implants were tested post-perfusion to confirm efficacy. On separate manipulation days, laser light was delivered as a 0.5 sec pulse gated at the start of a trial, a 0.5 sec pulse upon arrival at the middle of the maze, or as a cycled pulse of 0.5 sec on/off for the full run duration. Days with light delivery at run-start or mid-run were counterbalanced across groups on both tasks, with cycled light delivery always occurring last. Maze behavior and task events (e.g., start time, trial end, etc.) were recorded by the Ethovision XT tracking software via an overhead digital camera connected to a computer. Accuracy of automated behavioral measurements were verified through videotaping and hand scoring of a subset of sessions. Errors due to poor tracking found in Ethovision data post-processing were corrected by these hand-scored analyses directly from individual trial videos. On rare occasions a mechanical issue would occur and such trials were omitted from analysis.

### Reward Devaluation

Free access to the task reward (~2.5 g = ~50 pellets) was given to rats for 30 min. in their home cage in a separate room from the maze. Afterwards, injections of lithium chloride (0.3 M, 10 ml/kg, i.p.) were given to introduce nausea. Three of these pairings were given at 48 hr intervals. Reward pellet consumption was recorded prior to and after each procedure. Pellet consumption during task performance was also recorded before and after the devaluation procedure.

### Histology

At the termination of each experiment, a lethal dose of anesthesia (sodium pentobarbital) was administered, followed by a transcardinal perfusion of 0.9% saline and 4% paraformaldehyde. Brains were put in 20% sucrose solution overnight and frozen at −80 °C. Brains were sectioned under a microtome at 30-60 μm, mounted to slides, and cover slipped with a DAPI-containing medium. Fiber placement and zones of EYFP expressing neurons were assessed using fluorescent microscope analysis (Olympus BX53 fluorescent microscope with DP73 camera). For estimates of the efficacy of viral infection, brain sections were immunostained to label EYFP (primary and secondary antibodies: Rabbit Anti-GFP/Alexa 488 Goat Anti-Rabbi; Mouse Anti-eYFP/Alexa 594 Goat Anti-Mouse) and mounted to slides with Neurotrace to label neurons. Sections were then analyzed using a confocal microscope (Zeiss LSM880 with Airyscan laser scanning confocal microscope; Dartmouth College Biology Department Light Microscopy Core Facility). Image acquisition and analysis was quantified using Lmaris and Zen Blue. EYFP, neurotrace, and double-labeled neurons were separately counted in pseudorandom sections of DLS using a grid approach from three brains.

### Spiking data acquisition and processing

Subjects (n = 3) were given ChR2-containing viral construct injections as above and implanted with an optical fiber (200μm core, 0.39 NA) surrounded by eight independently moveable tetrodes (“VersaDrive-8 Optical”, Neuralynx) in either a chronic (n = 2; recordings during free behavior) or an acute preparation (n = 1; recordings under urethane anesthesia at 1.5g/kg). TTL timestamps from the laser control system and spiking activity (filtered at 600-6000 Hz) were collected using a Digital Lynx SX acquisition system and a pair of HS-18-MM preamplifiers (Neuralynx) with all signals referenced to a skull screw above the cerebellum. During recordings, cycles of blue light deliveries (2-5 mW) were given at 0.5 sec pulse duration every min, or 0.5 sec on/off pulses, in order to approximate task conditions. Single sessions included both light delivery protocols when recording stability allowed. Between sessions, tetrodes were lowered and the acquisition of new units confirmed visually. Units were isolated offline (Offline Sorter, Plexon), plotted (Supplementary Fig. 1), and analyzed for responsivity to light delivery (NeuroExplorer, SPSS). A unit was considered responsive if spiking frequency during the light delivery period went beyond a 95% confidence interval of baseline spiking for ≥ 3 consecutive 0.02 sec time bins. Unit activity was clearly distinguishable from photoelectric artifacts.

### Statistical analyses of behavior

Repeated measures ANOVAs were used to compare differences of dependent behavioral variables (e.g., cumulative maze duration, deliberations (VTEs) at choice point, accuracy, etc.) for within-subject and between-subjects analyses for main effects of the following factors: illumination day (e.g., baseline, run-start, mid-run, cycled on/off), trial within sessions, virus group (e.g., control vs. ChR2 vs. eNpHR3.0), and their interactions. These comparisons were conducted separately for four stages based on *a priori* events of interest: (1) task acquisition; (2) illumination tests after acquisition; (3) across pre- and post-devaluation extinction probe days; (4) during the reacquisition day (the rewards consumed measure was also compared to an equivalent pre-devaluation illumination day to confirm devaluation in the task). If there was a significant main effect for a behavioral variable or interaction between variables for either of the maze tasks (in addition to the post-devaluation reacquisition probe day in both), we used a Tukey-corrected post-hoc analysis for individual comparisons and individual day comparisons using univariate ANOVAs.

Contributions
KSS and ACGC: designed research, analyzed data, wrote the paper; ACGC, FS, AGM, MVDM, and EC: Performed research.

## Acknowledgements

We thank Kenneth Amaya, Jacqueline Perron-Smith, Elizabeth McNalley, Alyssa DiLeo, Alex Brown, and Dr. Stephen Chang for assistance. This work was supported by an NSF research grant to KSS (IOS 1557987).

